# Complex interplay between gene deletions and the environment uncovers cellular roles for genes of unknown function in *Escherichia coli*

**DOI:** 10.1101/2025.02.11.637708

**Authors:** Kaat Sondervorst, Kristina Nesporova, Matthew Herdman, Bart Steemans, Joëlle Rosseels, Sander K. Govers

**Author notes:** Address correspondence to Sander K. Govers,.

## Abstract

Phenotypic outcomes can be heavily affected by environmental factors. In this study, we exploited the previously observed nutrient-dependency of cell biological phenotypic features, captured by a cross-condition image-based profiling assay of *Escherichia coli* deletion strains, to examine this in more detail. We identified several general principles, including the existence of a spectrum of deviating phenotypes across nutrient conditions (i.e., from nutrient- or feature-specific to pleiotropic phenotypic deviations), limited conservation of phenotypic deviations across nutrient conditions (i.e., limited phenotypic robustness), and a subset of nutrient-independent phenotypic deviations (indicative of consistent genetic determinants of specific phenotypic features). In a subsequent step, we used this cross-condition dataset to identify five genes of unknown function of which the deletion displayed either nutrient-independent phenotypic deviations or phenotypic similarities to genes of known function: *yibN*, *yaaY*, *yfaQ*, *ybiJ*, and *yijD*. These genes showed different levels of phylogenetic conservation, ranging from conserved across the tree of life (*yibN*) to only present in some genera of the Enterobacterales (*yaaY*). Analysis of the structural properties of the proteins encoded by these y-genes, identification of structural similarities to other proteins, and the examination of their subcellular localization yielded new insights into their contribution to *E. coli* cell morphogenesis, cell cycle progression and cell growth. Together, our approach showcases how bacterial image-based profiling assays and datasets can serve as a gateway to reveal the function of uncharacterized proteins.

**Importance:** Despite unprecedented access to genomic information, predicting phenotypes based on genotypes remains notoriously difficult. One major confounding factor is the environment and its ability to modulate phenotypic outcomes. Another is the fact that a large fraction of protein-coding genes in bacterial genomes remains uncharacterized and have no known function. In this work, we use a large-scale cross-condition image-based profiling dataset to characterize nutrient-dependent phenotypic variability of *E. coli* deletion strains and exploit it to provide insight into the cellular role of genes of unknown function. Through our analysis, we identified five genes of unknown function that we subsequently further characterized by examining their phylogenetic conservation, predicted structural properties and similarities, and their intracellular localization. Combined, this approach highlights the potential of cross-condition image-based profiling, which extracts many cell biological phenotypic readouts across multiple conditions, to better understand nutrient-dependent phenotypic variability and uncover protein function.

## Introduction

A cellular phenotype is determined by the interaction of a cell’s genotype with stochastic, epigenetic and environmental factors (1–4). Uncovering the intricate relationships between these aspects is crucial in understanding the emergence of different cellular phenotypes and establishing clear genotype-phenotype associations. A phenotype itself is also complex and multifactorial, consisting of many, often interdependent traits, ranging from cellular fitness and growth capacity to cell morphological, intracellular organization and cell cycle aspects. In model microbes such as yeast and *E. coli,* systematic genome-wide screens, made possible through the construction of arrayed knockout collections, transposon mutant libraries or CRISPR-based approaches (5–11), have enabled the identification of genetic factors that contribute to specific phenotypic traits. For example, such screens have identified genes required for swarming motility (12), for growth across different environments (13), and for T7 bacteriophage growth (14) in *E. coli*.

While these initial screens typically focused on specific phenotypic traits, the advent of high-content screening methods has enabled the simultaneous readout of many phenotypic parameters. One prime example is image-based profiling, in which a large number of quantitative cellular and subcellular features are extracted from microscopy images of individual cells (15–18). This approach captures high-content cell biological information and provides more detailed insights into cellular physiology and the many aspects that constitute a given phenotype. Image-based profiling approaches across the genome-wide deletion collection of *E. coli* have revealed that a large fraction of the non-essential genome (∼20%) affects different aspects of cell morphogenesis, cell cycle progression, and growth (19–21). However, these screens were performed in a single environment, and we have recently shown that a large fraction of this genomic commitment is in fact nutrient-dependent (22). More specifically, we found that the majority of deviating phenotypes (∼75%), for all extracted phenotypic features, exhibited nutrient-dependency and did not deviate consistently across nutrient conditions. In line with this, a screen of the *E. coli* single-gene deletion library across 30 different nutrient conditions highlighted variable growth phenotypes across conditions for many deletion strains (23). These findings underscore the important role of the environment (in this case, the nutrients available in the growth medium) in determining phenotypic outcomes.

The goal of this study was to provide more detailed insights into the nature of this nutrient-dependency of phenotypic landscapes, while also exploiting both the extent (i.e., the many phenotypic features that were extracted) and the nutrient-dependency of these landscapes to uncover potential cellular roles for genes of unknown function. Even in a well-studied model system as *E. coli*, about one third of genes lack experimental evidence of function (24, 25). These unannotated genes are typically referred to as ‘y-genes’, stemming from their primary names starting with the letter ‘y’ (26). As metagenomic approaches have provided us with unprecedented access to bacterial genome sequences and the genes that lie within them, such sequence-to-function gaps are becoming increasingly more common in microbiology (27–29). Addressing this gap is challenging given the relatively slow process of experimental characterization, yet it is crucial in expanding our understanding of genetic diversity and cellular functioning, and further optimizing computational gene function predictions. One way to assign a function to a gene relies on the investigation of the consequences of gene inactivation (e.g., through deletion or transposon insertion). In earlier genome-wide screens, this was achieved by examining the growth or fitness of specific strains in which the genes of interest have been deleted or inactivated, across different nutrient conditions (13, 28–30). Here, we extended this approach by examining cross-condition image-based profiles of *E. coli* strains in which such y-genes were deleted (22). We exploited the high-content phenotypic characterization across conditions to identify specific y-gene deletion strains with nutrient-independent deviations of phenotypic features or phenotypic similarities to genes of known function. This yielded a set of five interesting candidates that were subjected to bioinformatic analyses (to assess their level of conservation), structural predictions (to identify their structural properties and homologies with other proteins), and fluorescent labeling (to determine their intracellular localization). Together, our experiments reveal crucial insights into the cellular role of these uncharacterized proteins and their contribution to the cellular functioning of *E. coli*.

## Results

### Re-analysis of cross-condition image-based profiling data reveals specific and pleiotropic deviating phenotypes

To examine the apparent nutrient-dependency of phenotypic landscapes in more detail, we started from an existing dataset that we generated in a recent study (22). This dataset contains population-level cellular features, extracted from phase contrast and fluorescence microscopy snapshots, for 806 *E. coli* deletion strains. These strains were equipped with fluorescent fusions to a cell division protein (FtsZ) and a DNA replication marker (SeqA), and sampled across four nutrient conditions (Fig. S1A). The four nutrient conditions varied substantially in carbon source and nutrient quality, and were based on M9 buffer, supplemented with either L-alanine (M9Lala), glycerol (M9gly), glucose (M9glu), or L-arabinose with casamino acids and thiamine (M9LaraCAAT). While the original dataset consisted of 77 average population-level features (related to cell morphology, nucleoid morphology, divisome formation, DNA replication, and growth) for each deletion strain per nutrient condition (22), we expanded this dataset for the current study to 96 features (Fig. S1B). The additional features consist of coefficient of variations (CVs) for specific cellular properties, which provide a measure for the variability of that property and thus also the level of control that is exercised over that property in a given nutrient condition. For all features in a given nutrient condition, we also calculated normalized scores (*s*), which represent the extent to which a mutant strain deviates from its respective wildtype for that feature and nutrient condition (see Materials and Methods). Typically, a threshold score |*s*| ≥ 3 for a specific feature is used to designate a deletion strain as one with a significant deviation or defect for that feature. A general overview of the distribution of deviating scores per feature and nutrient condition is provided in Fig. S1B. The phenoprint of a deletion strain is given by the collection of all its feature values and scores across conditions. In total, each phenoprint of our current dataset consists of 384 features (96 per nutrient condition for each strain). Both the raw measurements and normalized scores, for all strains across four nutrient conditions, can be found in Dataset S1.

In a first step, we investigated the specificity of deviating phenotypes for a given strain by capturing the extent to which a deviation is restricted to a given feature or nutrient condition. For this, we quantified both the number of nutrient conditions in which a strain displays a deviation (i.e., number of nutrient conditions with |*s*| ≥ 3 for at least one feature), as well as the number of features for which a strain displays a deviation (i.e., number of features with |*s*| ≥ 3 in at least one nutrient condition) (Fig. 1A). This analysis revealed the existence of a broad range of deviating phenotypes, with strains displaying very specific phenotypes (e.g., deviating in only a single feature in a single nutrient condition), more pleiotropic phenotypes (e.g., deviating in >90 features across all four nutrient conditions), and everything in between (Fig. 1A). Together, this spectrum of deviating phenotypes illustrates the heterogeneity in phenotypic alterations across nutrient conditions.

**Figure 1.**
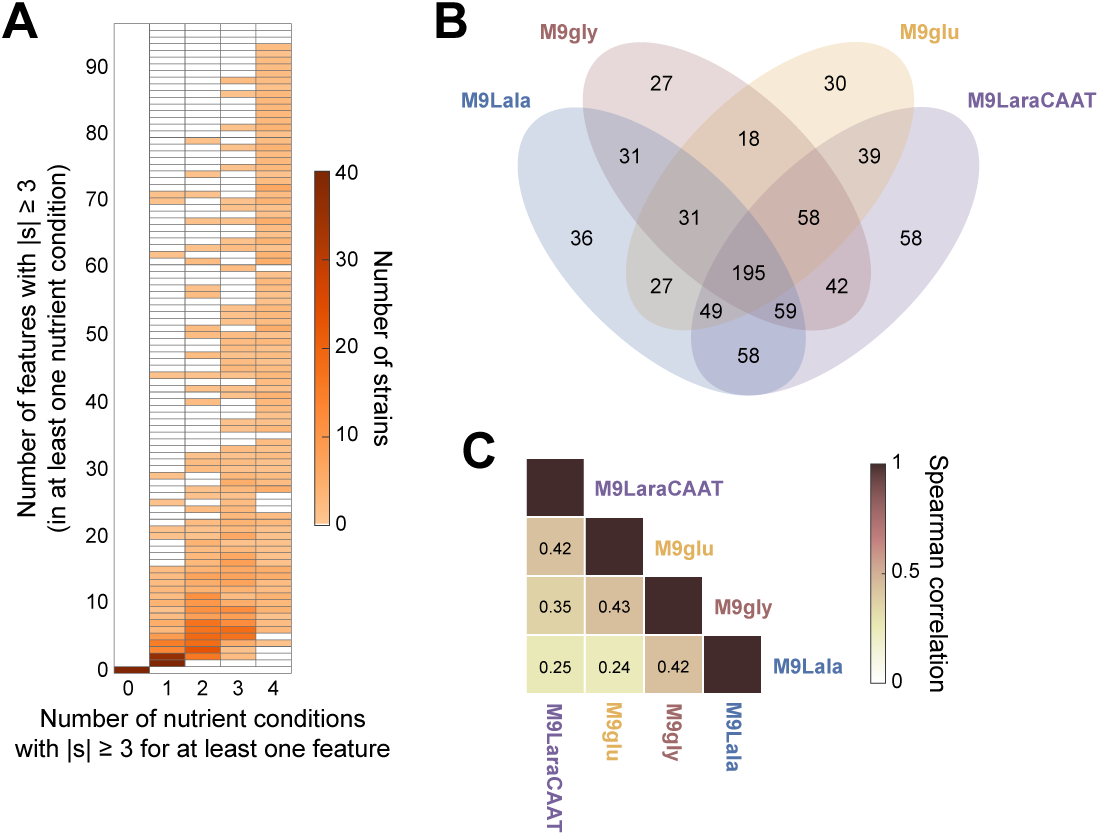
Diversity of deviating phenotypes and limited phenotypic robustness of deviations across nutrient conditions. A. Heatmap showing the distribution of deletion strains upon quantification of the number of features that deviate (|s| ≥ 3) in at least one nutrient condition versus the number of nutrient conditions in which at least one feature deviates. B. Four-way Venn diagram showing the number of strains that display a deviating phenotype (|*s*| ≥ 3 for at least one feature) across different combinations of nutrient conditions. C. Heatmap showing average correlations of deviating phenotypes across nutrient conditions. For each feature, only deletion strains displaying a deviating phenotype (|*s*| ≥ 3) in at least one nutrient condition were considered upon calculation of the correlation coefficients, which were then averaged across features.

### Limited robustness of phenotypic deviations across nutrient conditions

To probe how this heterogeneity affected the conservation of phenotypes of specific mutants across different nutrient conditions, we first examined in which nutrient condition strains displayed a deviating phenotype (|*s*| ≥ 3 for at least one feature). We found that 195 strains (24.2% of all deletion strains) displayed a deviation in all four nutrient conditions, whereas 197 (24.4%), 215 (26.7%), and 151 (18.7%) strains did so in three, two and one out of four nutrient conditions, respectively (Fig. 1B). In addition, we detected an overlap in deletion strains with deviating phenotypes between all possible combinations of nutrient conditions, indicating extensive nutrient-dependent phenotypic variability. This extensive phenotypic variability resulted in limited correlations of phenotypic scores of strains across conditions (Fig. 1C). For these correlations, only deletion strains with a deviation (|*s*| ≥ 3) in at least a single nutrient condition were considered per feature, and correlations were then averaged across features. While comparisons between more similar nutrient conditions (in terms of richness, as measured by the growth rates and cell sizes they support) gave rise to higher correlation values than more dissimilar nutrient conditions (Fig. S1C), the overall correlation values remained low, indicating limited robustness of deviating phenotypes across nutrient conditions in general.

### Nutrient-independent phenotypes enable the identification of genetic determinants of cell morphogenesis and cell cycle progression

While the preservation of phenotypes was generally limited across nutrient conditions, we were able to identify a number of nutrient-independent outliers, defined as having s > 3 (or s < −3) for at least one nutrient condition and s > 1.5 (or s < −1.5) in all other conditions, for almost all extracted features (Fig. 2A). The majority of strains displaying a nutrient-independent deviation did so for a limited number of features (Fig. 2B), although we also found some pleiotropic phenotypes (with >40 altered features per strain) that were nutrient-independent (Fig. 2B). The nutrient-independent nature of the deviations enabled us to more confidently identify genetic determinants of specific cellular features (Fig. 2C). An overview of all nutrient-independent outliers for all features is included in Dataset S1. Similar to our previous study that focused on cell cycle laws (22), many of the genes deleted in the strains that display nutrient-independent deviations encode proteins that localize to either the cell envelope or the nucleoid (Fig. 2C). Most noteworthy was the observation that almost all deletion strains displaying a consistently increased cell width were deletions of cell envelope components (17/22 or 77.3%; Fig. 2C), indicating an important link between envelope integrity and the determination of cell width.

**Figure 2.**
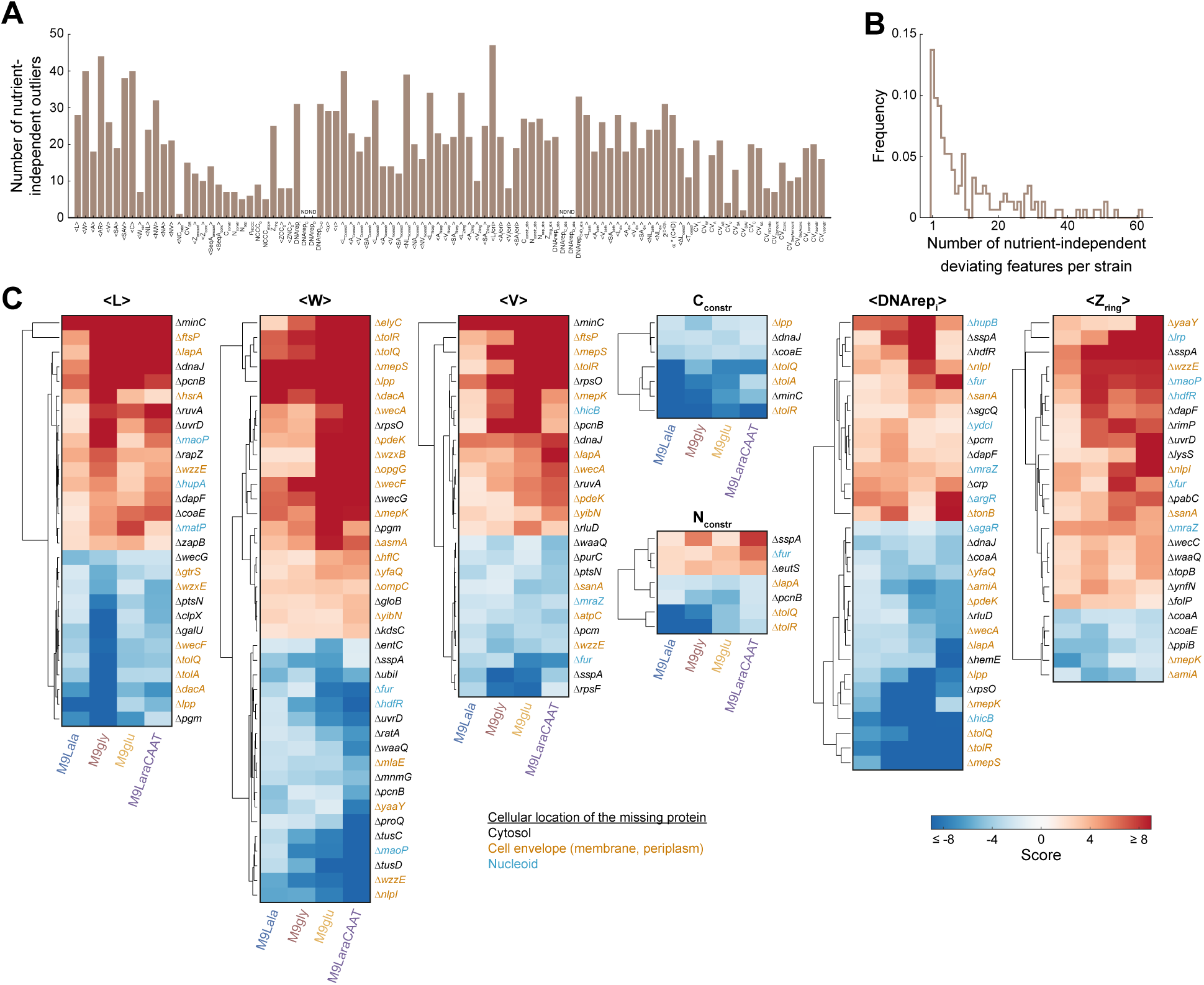
Identification of the genetic determinants of cell morphogenesis and cell cycle progression using nutrient-independent deviations of deletion strains. A. Bar graph showing the number of nutrient-independent outliers for each feature. Nutrient-independent outliers were defined as having s > 3 (or s < −3) for at least one nutrient condition and s > 1.5 (or s < −1.5) in all other conditions. B. Frequency distribution of the number of nutrient-independent deviating features per strain. C. Clustergrams showing the normalized scores of nutrient-independent outliers for the indicated features.

From this analysis, two y-gene deletion strains with interesting nutrient-independent phenotypes also emerged (Fig. 2C). A first one was the Δ*yibN* strain, which consistently gave rise to wider cells across conditions (Fig. 2C). Similar to other deletion strains that gave rise to wider cells, YibN contains a predicted transmembrane region and is expected to localize to the inner membrane. In addition, this strain also displayed longer cells in richer nutrient conditions, giving rise to markedly larger cells (in terms of cell volume and surface area) in these conditions (Fig. S2A). Another remarkable y-gene deletion strain was the Δ*yaaY* strain, which displayed more narrow cells and a later relative timing of FtsZ ring formation across nutrient conditions (Fig. 2C). In the richest nutrient condition, this strain also displayed a more pleiotropic phenotype, with cells being smaller on average (in terms of both length and width) with a large number of additional phenotypic alterations (Fig. S2A). This observation could be a consequence of significant growth defects, illustrated by our inability to obtain reliable growth measurements for this strain across all nutrient conditions (Fig. S2A). Together, these observations implicate both y-genes in important aspects of cell morphogenesis, cell cycle progression, and growth. As such, these mutants were retained for further analysis and characterization, after identification of additional y-gene deletion strains with relevant nutrient-dependent phenotypes in the subsequent section.

### Cross-condition phenotypic profiling can inform on cellular roles of genes of unknown function

An additional advantage of phenotypic profiling across multiple nutrient conditions is that it allows the identification of deletion strains with similar phenoprints in a more reliable and non-nutrient-specific way (i.e., even if certain phenotypic alterations are not preserved across nutrient conditions). We achieved this by examining correlations between the obtained multi-condition phenoprints of strains. High phenoprint correlations between two deletion strains, both positive and negative, are predictive of functional connections between the deleted genes and can inform on gene function (13, 31). To identify deletion strains with related phenoprints, we first calculated pairwise correlation scores for all strains that were sampled across all 4 nutrient conditions (Fig. 3A). This yielded a broad distribution of correlation coefficients (Fig. 3B), of which some of the highest absolute values were obtained between strains in which genes were deleted that encode proteins known to be involved in similar cellular processes (Fig. 3C), serving as a proof-of-concept.

**Figure 3.**
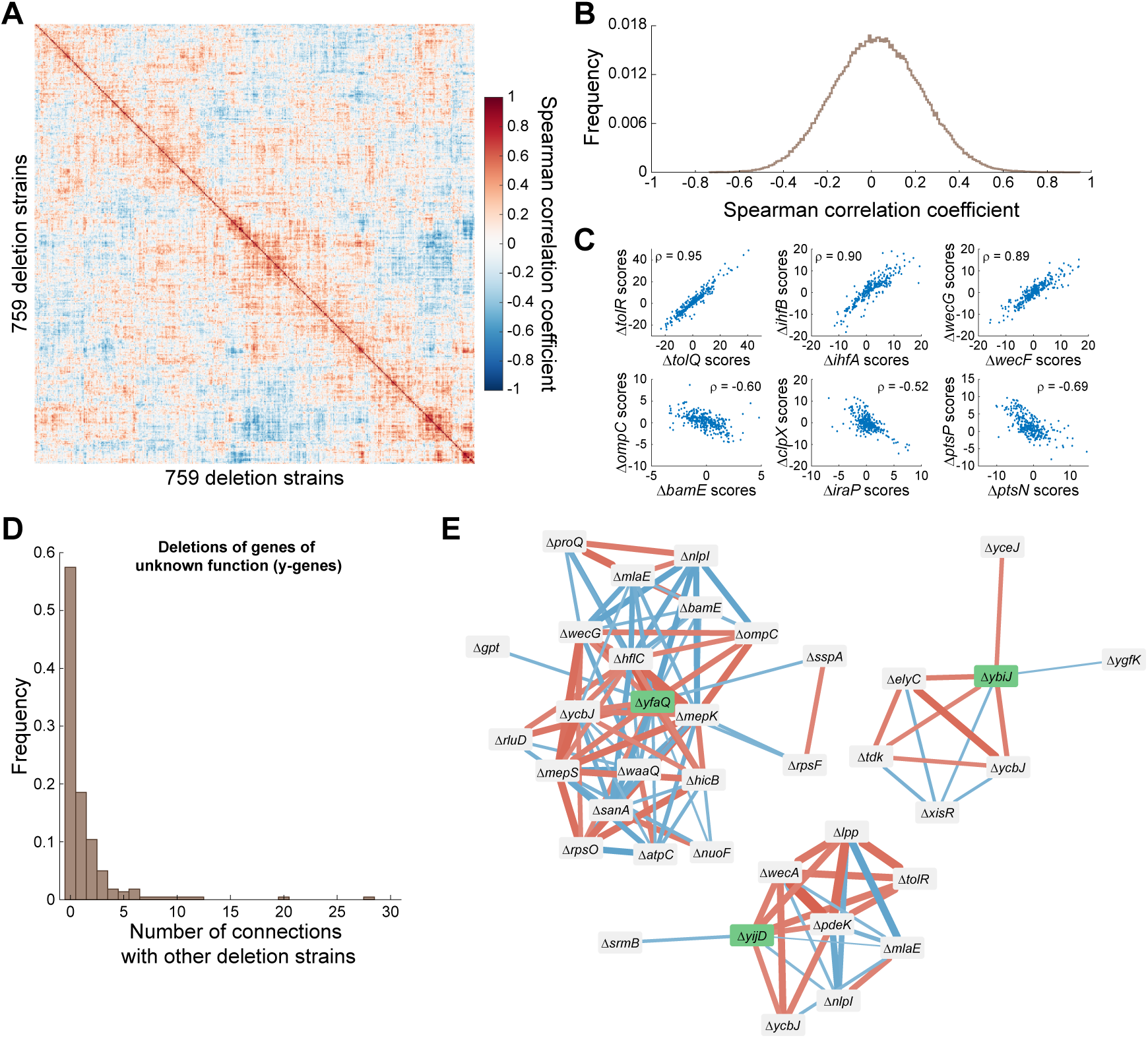
Phenoprint correlation networks of deletion strains can inform on cellular roles of deleted genes. A. Heatmap showing the correlations between cross-condition phenoprints of 759 deletion strains. B. Frequency distribution of all pairwise phenoprint correlations C. Examples of strong positive (top row) and negative (bottom row) correlations between phenoprint scores of strains in which the deleted gene products are known to be involved in similar or opposing cellular functions. D. Frequency distribution of the number of significant phenoprint connections between strains with deletions of y-genes and other deletion strains. E. Network representation of significant phenoprint correlations between deletions of the indicated y-genes and other deletion strains. The color of the edges indicates the sign of the correlation (blue: negative, red: positive), their width the relative strength of the correlation.

For example, TolR and TolQ are both inner membrane components of the Tol-Pal system that spans the entire cell envelope and is crucial for maintaining its integrity (32, 33). IhfA and IhfB are the two subunits of the nucleoid-associated protein IHF that specifically binds DNA, introduces sharp DNA bends, and affects gene expression (34–36). WecG and WecF both catalyze cytoplasmic steps in the biosynthesis of enterobacterial common antigen (37). For all these pairs, their positive phenoprint correlation exemplifies their similar roles in the same cellular process or pathway (Fig. 3C, top row). In contrast, strong negative correlations often indicated opposing roles (Fig. 3C, bottom row). For example, BamE is a lipoprotein that is part of the β-barrel assembly machinery, the Bam complex, which plays a crucial role in the biogenesis of integral outer membrane proteins in Gram-negative bacteria (38–40). Deletion of *bamE* leads to jamming of the Bam complex by RcsF, a Bam complex substrate and outer membrane protein-dependent surface-exposed lipoprotein involved in stress sensing (41–43). Surface exposure of RcsF normally occurs via the assembly of complexes with various outer membrane proteins, including OmpC (44, 45). Both BamE and OmpC are thus functionally connected to RcsF, but their deletion leads to different RcsF-related outcomes (i.e., jammed in the Bam complex vs. not in complex with OmpC) that likely underlie the contrasting phenotypes of their respective deletion strains. In other cases, the opposite roles are more clear, as IraP is an anti-adaptor protein of the ClpXP protease (46, 47), and PtsN and PtsP are competing parts of the nitrogen-related phosphotransferase system of *E. coli* (48, 49).

In a subsequent step, we exploited this approach to provide more insight into the cellular role of specific y-genes by examining potential connections of their deletion phenoprint with that of other genes of known function. For each y-gene deletion strain, we identified significant connections by using empirical thresholds that exclude the middle 99.7% of all correlation coefficients (Fig. 3D). This analysis showed that close to 50% of y-gene deletion phenoprints displayed significant connections to other deletion phenoprints, ranging from just one to many significant connections (Fig. 3D). This distribution of significant connections was similar to that obtained when examining the phenoprint connections of all deletion strains (Fig. S2B). An example of a y-gene deletion strain with many connections was the Δ*yfaQ* strain, which displayed connections to several deletion strains of genes involved in peptidoglycan or membrane remodeling and integrity (Δ*mepK*, Δ*mepS*, Δ*wecG*, Δ*ompC*, Δ*bamE*, Δ*mlaE*, Δ*nlpI*, Δ*waaQ*, Δ*sanA,* Δ*hflC*; Fig. 3E). Together with its predicted signal sequence and phenoprint (wider cells across nutrient conditions; Fig. S2A), this connection network suggested a potential role for the YfaQ protein in cell envelope assembly and/or structuring. Other y-gene deletions that displayed interesting connections and phenoprints included the Δ*ybiJ* strain, which had positive connections with the Δ*elyC* and Δ*ycbJ* strains (Fig. 3E) while displaying smaller and larger cells in nutrient-poor and nutrient-rich conditions, respectively (Fig. S2A). ElyC is an envelope biogenesis factor (50), whereas the gene encoding YcbJ overlaps with that of ElyC (opposite orientation), leading to the inactivation of *elyC* in a *ycbJ* deletion mutant. YbiJ contains a signal sequence, suggesting it localizes to the cell envelope and could also be involved in maintaining its integrity. Another notable y-gene deletion strain was Δ*yijD*, which displayed gradually wider cells in increasingly richer nutrient conditions, a slower growth rate in nutrient-poor conditions, early cell and nucleoid constriction across all conditions, and positive connections to Δ*tolR* and Δ*lpp* (Fig. 3E and S2A). As both these proteins are involved in linking different parts of the cell envelope or are parts of complexes that do so (33, 51), YijD, shown to be an inner membrane protein (52), could play a similar role or be part of a similar complex.

### Differences in phylogenetic conservation of identified genes of unknown function

To investigate the level of evolutionary conservation of the five identified y-genes, we first evaluated the occurrence of orthologs across the tree of life (Fig. 4A and S3). Of our five genes, *yaaY* displayed the most limited conservation (Fig. 4A-B), and was only found in a subset of genera of the Enterobacterales (i.e., *Escherichia*, *Salmonella*, and *Citrobacter*). While *yijD* was mostly conserved across the Enterobacterales, it was also present in most Vibrionales (Fig. 4A-B), including in notorious pathogens such as *Vibrio cholerae*. This contrasts with *ybiJ*, which was found across Enterobacterales, but not in other Orders (Fig. 4B and S4). Both *yfaQ* and *yibN* displayed notable conservation patterns. While *yfaQ* exhibited a sparse pattern across the bacterial domain, *yibN* orthologs were found across the entire tree of life (Fig. 4A-B), even in archaea and eukaryotes.

**Figure 4.**
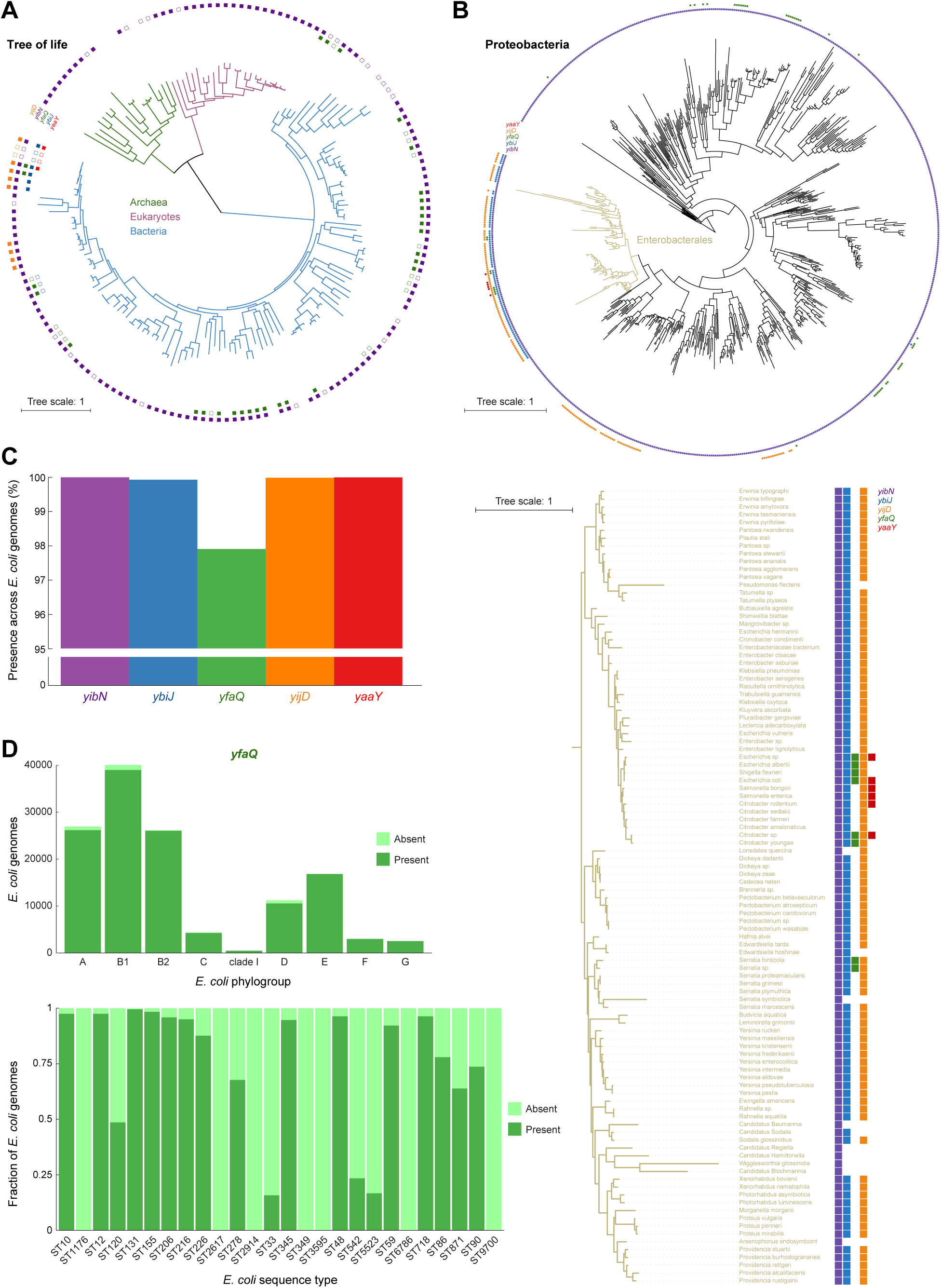
Conservation analysis of selected y-genes. A. The presence of the indicated y-gene orthologs, determined using the EggNOG database, was mapped onto the tree of life. Full squares indicate the presence of the respective ortholog in a specific species, while empty square indicate the presence of the ortholog in the relevant genus of the species present in the tree (but not in that species itself). The main tree branches represent the domains of Archaea (green), Eukaryotes (purple) and Bacteria (blue). Species names can be found in Fig. S3. B. The presence of the indicated y-gene orthologs was mapped to the tree of YibN orthologs in Proteobacteria. The Enterobacterales branch (yellow) is depicted separately in detail including species names. The species names for the full tree can be found in Fig. S4. C. Bar graph indicating the prevalence of the y-genes across a large-scale genomic collection of *E. coli* (consisting of 131610 genomes). D. Patterns of presence or absence of *yfaQ* across *E. coli* phylogenetic groups (top) and in 26 sequence types (relative distribution) that had at least 10 genomes in which *yfaQ* was absent (bottom).

In a second stage, we examined the conservation of these five y-genes across a large-scale *E. coli* genome collection, which we recently assembled (consisting of 131610 *E. coli* genomes (53)). This analysis revealed that four genes (*yaaY*, *ybiJ*, *yibN* and *yijD*) were strongly conserved across *E. coli* strains, as they could be found in (almost) all genomes (Fig. 4C). The only exception was *yfaQ*, which was still present in approx. 98% of the genomes, but was notably depleted or even completely absent in strains of specific sequence types (Fig. 4D and Table S1). Together, this analysis revealed clear differences in the broadness of the phylogenetic spectrum in which the y-genes operate.

### Cell envelope-associated localization patterns of YibN and YbiJ

To provide further insight into the cellular role of these y-genes, we determined their homology (Table S2), performed structural predictions, and examined their intracellular localization. For this last step, we attempted to construct C-terminal msfGFP fusions to each of them at their native chromosomal locus. For all selected y-genes but *yaaY*, we successfully generated such a fluorescent fusion and subjected these strains to microscopy across the four different nutrient conditions of our initial screen (i.e., M9Lala, M9gly, M9glu, and M9LaraCAAT). While construction was successful for YfaQ-msfGFP, we could not detect a signal in cells producing this fusion across conditions. This could be a consequence of either low fusion protein levels in these conditions (y-genes often have lower expression levels (24)) or aberrant protein fusion production and/or functionality. YfaQ homology determination and structural predictions suggest a bipartite arrangement of the protein (Fig. S5 and Table S2), where the N-terminal domain shares features of transpeptidases, while the C-terminal domain is structurally similar to SpoIID transglycosylase (Fig. S5). YfaQ also harbours an N-terminal Sec signal peptide and SPI cleavage site, suggesting export to the periplasm (Fig. S5).

The three other remaining fusion proteins did display a distinct localization pattern. YibN-msfGFP displayed an envelope-associated localization across all conditions (Fig. 5A), which is in line with its predicted structure. YibN is predicted to consist of a short N-terminal transmembrane domain, formed by a hydrophobic, non-polar, α-helical tail, and a larger cytoplasmic globular domain that share homology with rhodanese domain proteins (Fig. 5B). Proteins harboring such rhodanese domains are ubiquitous across the tree of life (54), which likely explains the broad occurrence of *yibN* orthologs (Fig. 4A). Rhodanese domain proteins are implicated in a wide variety of cellular processes, mostly through their ability to bind and transfer sulfur atoms via a cysteine residue within their active site (54–56) (Fig. 5C-D). While the predicted catalytic domain of YibN contains such a canonical cysteine residue, it is on the opposite end in comparison to that of other rhodanese proteins (Fig. 5C-D).

**Figure 5.**
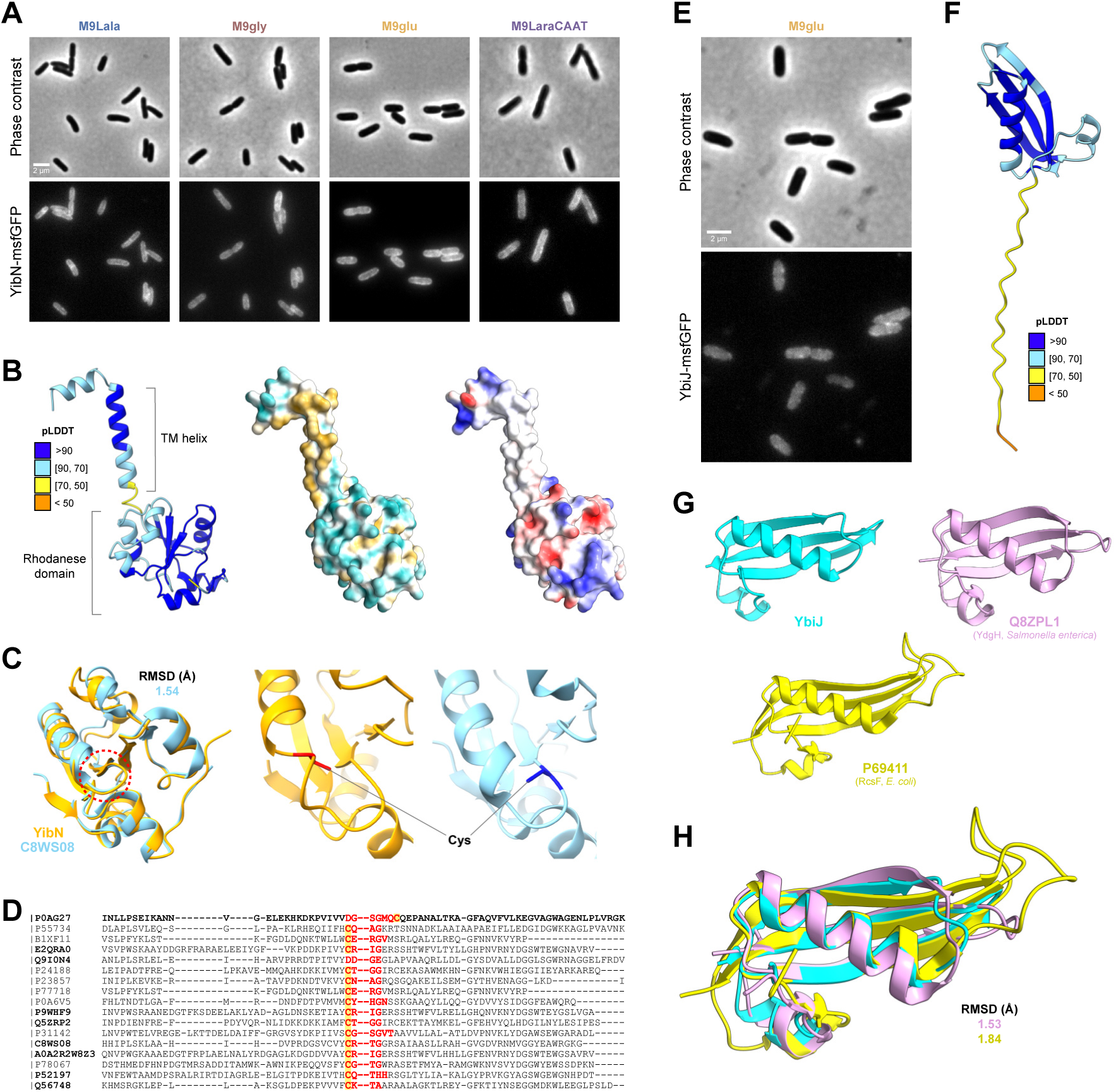
Localization and structural properties of YibN and YbiJ. A. Representative phase contrast (top row) and epifluorescence (bottom row) images of the MG1655 *yibN*::*yibN-msfgfp* strain across the indicated nutrient conditions. B. Structural depiction (ribbon) of YibN generated by AlphaFold 2 and coloured by pLDDT (left). Surface representations, coloured depending on hydrophobicity (middle) or electrostatic surface potential (right), are provided. C. Aligned rhodanese domains from YibN (AF2; orange) and C8WS08 (PDB 3TP9; blue) (left). The enzymatic loop has been indicated (red dashed circle). The cysteine residue and side chain are highlighted in red for YibN (middle) and blue for C8WS08 (right), showing that they lie on opposite ends of the loop. The low root mean square deviation (RMSD) indicates the good quality of the alignment. D. Aligned amino acid sequences of rhodanese domains. The first sequence is taken from YibN; other sequences depicted in bold were among the top 10 HHpred hits for YibN. The residues comprising the enzymatic loop for each protein have been highlighted in red; the putative enzymatic cysteine is highlighted in yellow. E. Representative phase contrast (top) and epifluorescence (bottom) images of the MG1655 *ybiJ*::*ybiJ-msfgfp* strain in M9glu. F. Structural depiction (ribbon) of YbiJ generated by AlphaFold 2 and coloured by pLDDT. The C-terminal peptide with pLDDT < 70 comprises the secretion signal and cleavage site. G. Structural depictions of the McbA-like domain of YbiJ (blue), YdgH (PDB 4EVU; pink) from *Salmonella enterica* and RcsF (PDB 6T1W; yellow) from *E. coli*. Structural homologies were determined based on HHpred results. H. Superimposed McbA-like domains showing structural homology. The low RMSDs indicate the good quality of the alignments.

YbiJ-msfGFP displayed a feint envelope-enrichment, but only in M9glu (as no signal could be detected in the three other nutrient conditions) (Fig. 5E). YbiJ is a small protein without notable predicted structural features, except that it contains a Sec secretion signal and likely localizes to the periplasm (Fig. 5F). YbiJ is a paralog of McbA, both containing a small domain of unknown function (DUF1471) (57). McbA is thought to be involved in the response to extracellular stress together with other DUF1471 proteins (57), although the exact role of these proteins remains unclear. Despite minimal sequence homology, YbiJ displays a distinct structural homology to RcsF (Fig. 5G-H and Table S2), the Bam complex substrate and outer membrane lipoprotein involved in envelope stress sensing (41, 42). YbiJ lacks the hydrophobic α-helical tail region of RcsF, suggesting it does not interact directly with the outer membrane. While the envelope-enriched localization pattern of the YbiJ-msfGFP fusion protein in M9glu is in line with the properties of YbiJ, we currently do not know why we did not observe this in other nutrient conditions, although nutrient condition-specific production of the protein could serve as a likely explanation.

### YijD localizes in envelope-associated clusters enriched at the cell subpoles and the division septum

The YijD-msfGFP fusion protein displayed a consistent, yet remarkable, localization across nutrient conditions (Fig. 6A). In each condition, YijD-msfGFP was clearly envelope-associated, in line with its reported inner membrane-association (52), but did so in a diverse number of irregularly shaped clusters that appeared to be enriched at the (sub)polar regions of the cells and the division septum (Fig. 6A). To provide further insight into this localization pattern, we constructed demographs, which consist of linear representations of integrated fluorescence of cells sorted by their cell length (58). These representations further supported an enrichment of YijD-msfGFP clusters at the subpolar regions (instead of the cell pole), which became more pronounced in rich conditions (Fig. 6B). Near the very end of the cell division cycle, clusters also emerged at the dividing septum (where new cell poles are formed after completion of cell division) (Fig. 6B). Examination of the predicted structure revealed properties in line with its membrane association, as YijD is predicted to be a small protein consisting of 4 hydrophobic and non-polar α-helices (Fig. 6C). Protein topology analysis revealed that helices can insert into and span the inner membrane (Fig. 6D). While these properties explain the observed envelope-associated localization, the mechanisms and reasons underlying the clustering and localization currently remain enigmatic.

**Figure 6.**
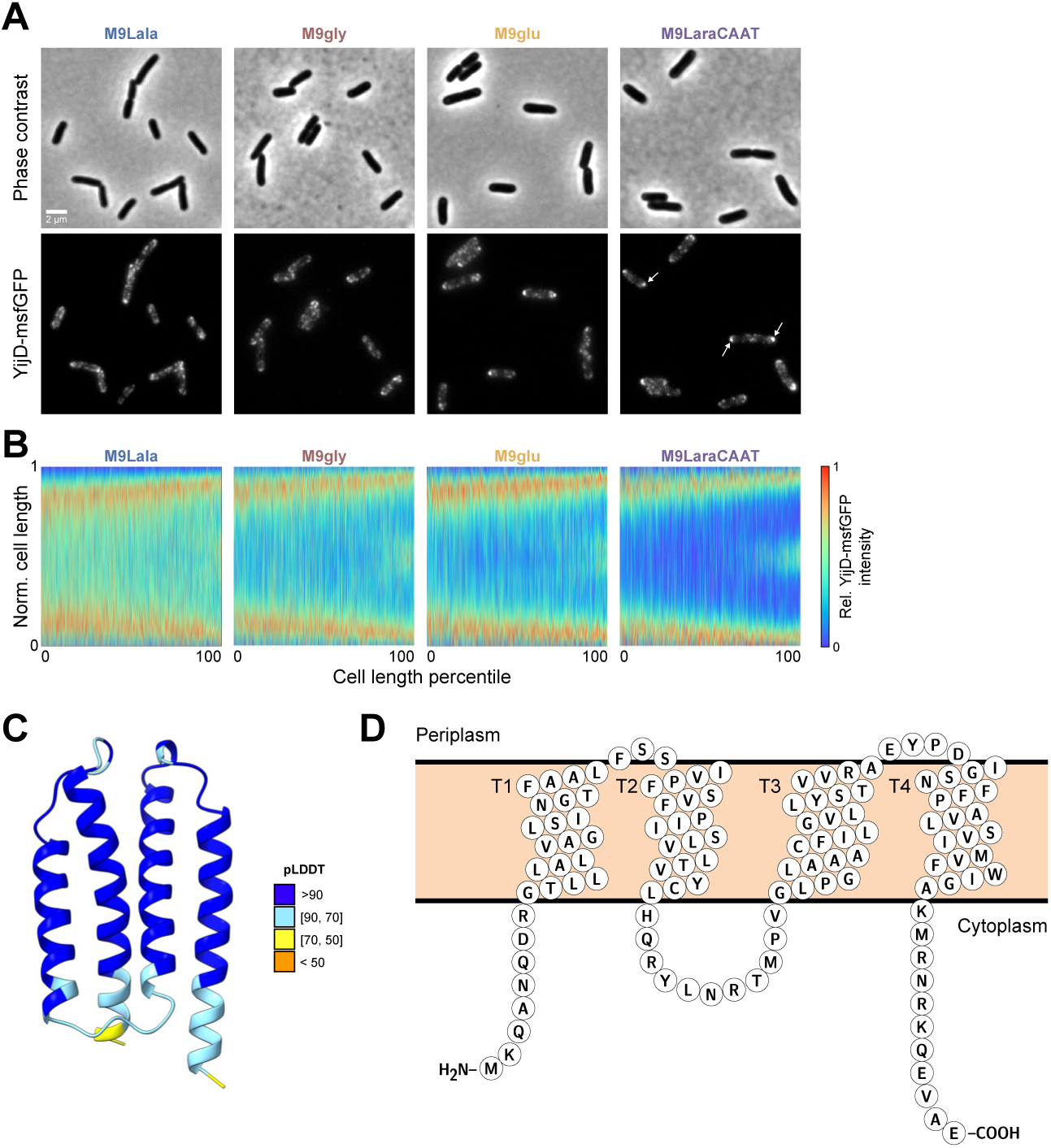
Localization and structural properties of YijD. A. Representative phase contrast (top row) and epifluorescence (bottom row) images of the MG1655 *yijD*::*yijD-msfgfp* strain across the indicated nutrient conditions. White arrows highlight subpolar localization of YijD-msfGFP clusters in M9LaraCAAT. B. Demographs of the YijD-msfGFP signal across the indicated nutrient conditions (n = 3997, 3121, 2125, and 2048 cells for M9Lala, M9gly, M9glu, and M9LaraCAAT, respectively). Cell length is normalized to show the relative cell length in the y axis. C. Structural representation (ribbon) of YijD protein generated using Alphafold2 and coloured by pLDDT. D. Protter representation of YijD homology illustrating transmembrane helices and cytoplasmic/periplasmic-exposed regions.

## Discussion

By further characterizing the extensive nutrient-dependency of deviating phenotypes of deletion strains, we were able to examine the nutrient-dependency of phenotypic landscapes and identify several general principles. A first was that a spectrum of deviating phenotypes exists across nutrient conditions, ranging from nutrient- and/or feature-specific, to more pleiotropic, with multiple phenotypic features being affected across nutrient conditions (Fig. 1A). A second was that phenotypic robustness was very limited across conditions, with the majority of deviating phenotypes not persisting across nutrient conditions (Fig. 1C). However, nutrient-independent phenotypes do exist and can be exploited to identify true genetic determinants of specific phenotypic features (Fig. 2). At the same time, we showed that similarities or dissimilarities in nutrient-dependent phenotypes of deletion strains, quantified by the correlation of their phenoprints, can be used to identify functional connections between genes and inform on their cellular function (Fig. 3). As such, microscopy-based phenoprint correlations provide an additional type of metric that enables investigation of protein interactions and function, complementary to existing databases (59, 60).

We further exemplified this by examining five specific genes of unknown function in *E. coli* that led to notable phenotypes upon their deletion. Of these five, *yaaY* remains the most enigmatic. While its deletion led to a pronounced cell morphological and cell cycle phenotype, with more narrow cells and a later timing of FtsZ ring formation across nutrient conditions (Fig. 2C), the gene displayed very little conservation outside of *E. coli*. In addition, we failed to construct a fluorescent protein fusion and its structural predictions yielded very poor results (Fig. S6). The protein has been computationally predicted to localize in the plasma membrane (61), yet this has not been experimentally verified. In contrast, we were able to experimentally verify the envelope-associated localization of the rhodanese domain-containing YibN (Fig. 5A). Structural predictions indicate it likely localizes there by inserting in the inner membrane using its N-terminal, hydrophobic, and non-polar α-helix (Fig. 5B). Recently, YibN has been identified as an interaction partner of YidC (62), modulating the latter’s ability to insert proteins into the inner membrane and organize its lipids (63–66). This interaction and regulating role of YibN is in line with the upregulation of YibN upon YidC depletion (67), and *yibN* being in an operon together with *grxC*, *secB*, and *gpsA*, all proteins with envelope-related functions (68). While *yibN* deletion did not lead to observable growth defects across conditions (Fig. S2A) or temperatures (62), the deletion strain did display consistently wider cells across conditions (Fig. 2C). This provides further support for the importance of envelope integrity in determining cell width and highlights the potential of image-based profiling for uncovering phenotypes that previously remained undetected. At the same time, the exact molecular mechanisms and cellular role of YibN, together with the part played by its rhodanese domain and active site with cysteine residue (Fig 5C), remain to be determined.

While YfaQ localization could not be experimentally verified using a fluorescent protein fusion, its predicted localization to the periplasm through a Sec signal peptide (Fig. S5), the phenoprint and connection network of its deletions strain (Fig. 3E and S2A), its homology determination (Table S2), and its structural predictions (Fig. S5) all point in the same direction: a role for the protein in maintaining envelope integrity by mediating peptidoglycan turnover or assembly, in line with its deletion leading to consistently wider cells (Fig. S2A). Another protein with a Sec signal sequence was YbiJ, although most notable insights into its cellular role came from its paralogues and structural homology determination that revealed a striking structural homology with RcsF (Fig 5E-H). Similar structural similarities between other DUF1471 proteins and RcsF have also been reported recently (69), and several of these proteins have been implicated in specific stress responses (70–72). In addition, expression of *ybiJ* was also shown to be upregulated under acid stress (73) and exposure to glutathione and ciprofloxacin (74), further hinting towards a role of this protein in stress sensing or mitigation. Although the precise contribution and molecular function of YbiJ and other DUF1471 proteins in these stress responses currently remains unclear, one intriguing hypothesis is that this could be mediated via the Rcs stress response, of which the activation is typically controlled by RcsF and its interaction with transmembrane outer membrane proteins (42).

YijD was the protein with the most distinct localization pattern, displaying a cluster-like organization that was enriched at the subpoles and the division septum (Fig 6A-C). While displaying predicted structural properties that match an inner membrane localization, it remains to be determined how the heterogenous and localized pattern is established. This could be an inherent property of the protein, or through interaction with other (membrane) proteins. The latter is in line with its phenoprint correlation network displaying connections to other proteins that are part of complexes that link different parts of the cell envelope (Fig. 3E). Independently of how YijD localization is established, it is yet another factor contributing to the inherent intra-cellular heterogeneity in the composition of membranes (75). Additional factors that have been demonstrated to display a heterogeneous membrane-associated localization pattern similar to that of YijD (i.e., preferentially located in the cell poles and septum) are cardiolipin and other anionic phospholipid domains (76, 77). Moreover, in its genomic context, *yijD* is located in an operon together with *fabR*, a negative regulator of unsaturated fatty acid biosynthesis (78–80). Together, this indicates a potential role for YijD in fatty acid and membrane homeostasis, although the exact nature of this remains to be elucidated.

In general, our ability to assess the contribution of genetic variation to specific phenotypic features has improved throughout the years (81, 82). Yet, predicting phenotypes using genotypic information remains challenging. This is due to pleiotropy, where a single genetic change affects two or more phenotypic traits (83), the fact that phenotypic traits are polygenic or even omnigenic, where many genes (indirectly) contribute to variations in many phenotypic features (84), and the potential of the environment to modulate phenotypes (e.g., the expression of antibiotic resistance genes in bacteria or colony morphology in yeast (85, 86)). In this study, we highlight the potential of cross-condition image-based profiling, where many phenotypic readouts are considered simultaneously across multiple conditions, to identify general principles of cross-condition phenotypic variability and serve as a gateway for uncovering protein function.

## Materials and methods

### Data analysis

For our analyses, we started from an existing dataset (22), which we expanded to 96 average population-level features by also including coefficient of variations (CVs) for specific cellular properties (obtained by dividing the standard deviation for a feature by its mean). For all 96 features in a given nutrient condition, we calculated normalized scores (s) as before (22): s = 1.35 x (F_i_ – median (F_i_^parent^))/iqr(F_i_^parent^). Here, F_i_ is the measured value of feature i in a nutrient condition, F_i_^parent^ are the values of the parent strain replicates (≥115 replicates) for feature i in the same nutrient condition, and iqr is the interquartile range. As the interquartile range of normally distributed data is 1.35 times their standard deviation, we scaled the normalized scores by this factor to express them in terms of standard deviations from the median of the parent strain values.

### Correlation coefficients

Spearman correlation coefficients (ρ) between phenoprints were calculated using MATLAB’s built-in corr function. To determine a cutoff for correlation significance, we first tried a permutation-based approach as in (13, 31) using 10000 permutations to construct a background distribution of correlations. As this approach yielded cutoffs that were not sufficiently stringent, we chose to identify significant correlations between the phenoprints of deletion strains using thresholds that exclude the middle 99.7% of all empirically obtained correlation coefficients: [-0.5154 0.6476]. For the generation of correlation networks, we used the built-in, edge-weighted, spring-embedded algorithm in Cytoscape (v3.10).

### Strains and growth conditions

Bacterial strains and primers used in this study are listed in Table S3 and S4, respectively. Strains were cultured in M9 medium, supplemented with 0.2% of L-alanine (M9Lala), glycerol (M9gly), glucose (M9glu), or L-arabinose, 0.1% casamino acids, 1 µg/ml thiamine (M9LaraCAAT). For microscopy, each strain was first grown to stationary phase in the appropriate nutrient condition in pre-culture tubes (3 ml of growth media) at 37°C under well-aerated conditions (200 rpm on an orbital shaker). A dilution (>1/10000) was then used to inoculate fresh growth medium (3 ml). The resulting cultures were allowed to grow until they reached an optical density at 600 nm (OD_600_) that corresponded to early exponential phase (OD_600_ = 0.10-0.15) before being sampled for microscopy.

### Construction of mutant strains

Mutant strains producing C-terminal translational fluorescent fusions to y-genes of interest were constructed by lambda Red-mediated recombination (87). Detailed construction procedures for each mutant strain are given in Table S3. All constructed mutants were initially confirmed by PCR with primer pairs attaching outside of the region where homologous recombination occurred (Table S4). Correct deletion or integration of PCR products was further verified by Sanger sequencing (LGC Genomics or Eurofins). Obtained reads were aligned to the template on Benchling using default parameters (Multisequence alignment, MAFFT v7), cleaned up, and checked for mutations.

### Microscopy

Cells were imaged on 1.5% agarose (UltraPure^TM^ Agarose, Invitrogen^TM^, Thermo Fisher Scientific, USA) pads, supplemented with M9 buffer. For each sample, 0.5 µl of cell culture was spotted on the pad, which was then covered with a cover glass (Fisherbrand™ Glass Square Coverslips, Fisher Scientific, Belgium). Per sample, 10 or 15 positions were imaged to ensure capture of a sufficient number of cells (n ≥ 2048 cells).

Imaging was performed on a Nikon Eclipse Ti2-E inverted epi-fluorescence microscope controlled by NIS-Elements AR software and equipped with a Plan Apo λ 100x DM oil objective (NA= 1.45; Nikon, the Netherlands), Ph3 condenser annulus (Nikon, the Netherlands), and a Kinetix A22J723021 sCMOS camera (Teledyne Photometrics, USA). For fluorescence excitation, a pE-800 light source (CoolLED, UK) with eight separate LED channels allowed narrowband excitation, and filter cubes with BrightLine® Pinkel filter sets (penta-band set: LED-DA/FI/TR/Cy5/Cy7-5X-A; Semrock, IDEX Health and Science, USA) and OptoSpin filter wheels (Cairn GmbH, Germany) allowed capturing of narrowband fluorescent emission signals. The following optical configuration was used to acquire fluorescent images: GFP (excitation (CoolLED): 470 nm, excitation filter: FF01-474/27, dichroic FF409/493/573/652/759-Di01, emitter filter cube: FF01-432/515/595/681/809, emitter filter wheel: FF01-515/30).

### Conservation analysis of y-genes

In order to explore the prevalence of the selected y-genes and their orthologs, we used the EggNOG (v.5.0.0) (88) function Sequence search (http://eggnog5.embl.de/#/app/seqscan), in which we uploaded the reference proteins from *E. coli* MG1655 (YbiJ WP_206053394.1; YijD WP_000806411.1; YfaQ MCK2194123.1; YaaY MCN6283483.1; YibN WP_001156181.1). We curated the EggNOG outcomes for each of these proteins to examine whether an ortholog is reported for a given species (or genus). After this, we mapped the information about ortholog presence/absence to phylogenetic trees using the Interactive Tree of Life tool (iTOL v6) (89). For this, we adjusted the tree of life provided by iTOL (https://itol.embl.de/itol.cgi) to not contain multiple representatives of the same species. To visualize the orthologs’ presence on a more nuanced level, we used another tree provided within EggNOG via ETE toolkit implementation. The tree was calculated for YibN orthologs on a taxonomic level of the phylum of Proteobacteria (syn. Pseudomonatoda). We filtered out cases of unspecific taxonomy (e.g., labeled only as alpha proteobacterium) and species that were represented by multiple strains to ensure only single species representatives. We chose the YibN-based tree as it was the most abundant of the examined proteins. A consequence of this is that the visualized phylogenetic relations between species can differ from trees based on more data than a single protein and that a YibN ortholog is automatically present in this visualization. Related to this, species associated with orthologs of one or more of the other four proteins, but not YibN, could have been omitted. The visualization in Fig. 4 does not depict the species name, yet this information can be found in Fig. S3-S4.

The within-species conservation in *E. coli* was done on the level of genes corresponding to the reference proteins. The genes were implemented into a database which was screened using ABRicate with a threshold of 85% for query coverage and identity against a large-scale *E. coli* genomic collection of 131610 genomes (53). The outcomes were further processed in RStudio (v2023.06.0) with R (v4.4.1) to merge them with the appropriate genome metadata, to summarize the results and to create figures. As *yfaQ* was the only gene that showed a slight decrease in prevalence among *E. coli* (i.e., present in ∼98% of genomes), we additionally examined its absence patterns on the level of *E. coli* sequence types (STs). From 332 different STs which had at least one genome with *yfaQ* absence, we focused on 26 STs which had minimally 10 genomes without *yfaQ* (Fig. 4D). Importantly, *yfaQ* absence is based on the chosen bioinformatic thresholds and does not completely exclude the possibility of these strains harboring a functional YfaQ or its orthologs.

### Homology and Structural Predictions

Sequence similarity searches and homology detections were performed with the MPI Bioinformatics Toolkit (90), using HHpred (91, 92) comparison methods and the y-gene protein sequences provided by UniProt. Searches were performed against the PDB_mmCIF70 (HMM-based search and alignment generation against PDB chains; filtered for a maximum sequence identity of 70%) or Pfam-A databases. For more accurate prediction of transmembrane domains and signal peptides, DeepTMHMM 1.0 (93) and SignalP 6.0 (94) were used. Schematic representation of YijD topology was generated using Protter (95).

Structural predictions of y-gene protein products were generated using AlphaFold2 (AF2) through the web version of ColabFold v1.5.5 (96). Query protein sequences were again taken from the UniProt database, as above. Predictions were run for a maximum of 12 recycles. Structural similarity of predicted protein structures with homologous protein structures (as identified by HHpred searches) was quantified using the pairwise structural alignment tool from RCSB PDB (97).

### Image analysis

Phase contrast images were processed using the ‘bact_phase_omni’ model from Omnipose (98). This model was employed to identify cell outlines. To eliminate incorrectly detected cells, a support vector machine model was trained to distinguish single cells from erroneous detections (99). This approach ensured that only single cells were included in the analysis, thereby minimizing potential biases. From the correctly identified cell outlines, demographs, which are linear representations of the integrated fluorescence signal across the population of cells, sorted by cell length, were constructed.

## Data and code availability

All datasets used in this manuscript are available in the supplemental information (Dataset S1). All original code has been deposited at https://github.com/Govers-Lab/Sondervorst_et_al_2025.

## Acknowledgements

We thank Dr. Christine Jacobs-Wagner for inspiring this work and Dr. Seung Hyun Cho for scientific insights and discussions related to the structural similarity of YbiJ with RcsF. We would also like to thank the members of the Govers laboratory for fruitful discussions and critical reading of the manuscript. K.S. was funded by doctoral fellowship from the Research Foundation – Flanders (file number 1110925N). K.N. was funded by the European Union under Horizon Europe: MSCA Actions (Project 101105027) and by a junior postdoctoral fellowship from the Research Foundation – Flanders (file number 1251624N). This work was also supported by a start-up grant (STG/21/068) from the KU Leuven Research Fund and by the European Union (ERC Starting Grant BacterialBlueprint 101040800).

